# Efficacy of Deferoxamine Mesylate in Serum and Serum-Free Media: Adult Schwann Cell Survival Following Hydrogen Peroxide Induced Cell Death

**DOI:** 10.1101/2025.01.23.634338

**Authors:** Yee H.E. Ma, Abhinay R. Putta, Cyrus H. H. Chan, Stephen Vidman, Paula Monje, Giles W. Plant

## Abstract

Schwann cell (SC) transplantation shows promise in treating spinal cord injury as a pro-regenerative agent to allow host endogenous neurons to bridge over the lesion. However, SC transplants face significant oxidative stress facilitated by ROS in the lesion leading to poor survival. Deferoxamine Mesylate (DFO) is a neuroprotection agent shown to reduce H_2_O_2_ induced cell death in serum containing conditions, here we show that DFO is not necessary to induce neuroprotection under serumfree conditions by cell survival quantification and phenotypic analysis via immunohistochemistry, Hif1a and collagen IV quantification via whole cell corrected total cell fluorescence, and cell death transcript changes via RT-qPCR. Our results indicate survival of SC regardless of DFO pretreatment in serum-free conditions and an increased survival facilitated by DFO in serum containing conditions. Furthermore, our results showed strong nuclear expression of Hif1a in serum-free conditions regardless of DFO pretreatment and a nuclear expression of Hif1a in DFO treated SCs in serum conditions. Transcriptomic analysis reveals upregulation of autophagy transcripts in SCs grown in serum-free media relative to SCs in serum conditions, with and without DFO and H_2_O_2_. Thus, indicating a pro-repair and regenerative state of the SCs in serum free conditions. Overall, results indicate the protectiveness of chemically defined medium in enhancing SC survival against ROS induced cell death *in vitro*.

## 1. Introduction

Spinal cord injury (SCI) is a severe, life-altering condition affecting over 500,000 individuals annually, predominantly resulting from preventable causes such as motor vehicle accidents and violence [1]. A hallmark of SCI is axon degeneration, where the synaptic connections between neurons are disrupted, leading to loss of function and further neurological complications [2–4]. Cellular transplantation therapies have emerged as a promising approach for axon regeneration, with autologous Schwann cell (SC) transplants currently being evaluated in clinical trials for their efficacy in treating cervical and thoracic SCI [5–14]. Originally proposed by David and Aguayo in 1981, peripheral nerve transplants were shown to promote regeneration within the central nervous system by grafting peripheral nerve tissues, which induced axonal regrowth in focal injuries [14]. Since then, SCs have been characterized as peripheral glial cells and have shown considerable regenerative potential in SCI models, with numerous preclinical studies highlighting their capability to facilitate axon regeneration [8–14]. Compared to other transplant paradigms such as neural progenitor cell (NPC) transplants [16,17], SCs offer several advantages. As SCs are mature cells, the risk of spontaneous differentiation into unintended cell types and the formation of ectopic colonies seen in NSC transplantations are eliminated [18]. Furthermore, the autologous nature of the transplants limits the need for immunosuppression following transplantation [11,19]. There are also standardized and straightforward methods to isolate and characterize SCs from a patient’s peripheral nerves, making them an accessible and consistent candidate for cellular transplantation therapy [20, 21].

Nonetheless, SC transplants face several challenges, particularly the acute death of transplanted cells caused by oxidative stress in the host spinal cord following injury. Reactive oxygen species (ROS) such as hydrogen peroxide, hydroxyl radicals, and superoxide anion radicals have been shown to induce oxidative stress and mediate cell death in spinal cord injury [22]. Thus, the efficacy of cellular transplant therapies has been limited due to the acute cell death that occurs post-transplantation. To address the issue, recent strategies has been focused on preconditioning SC with various pharmaceutical agents prior to transplantation [23–26].

One such agent of interest is Deferoxamine Mesylate (DFO) [22, 26]. Mechanistically, DFO acts as an iron (Fe^2+^) chelator, thus inhibiting the function of HIF-PHDs and other Fe^2+^, O_2_, and 2-oxogluterate dependent enzymes that mark HIF proteins for proteasomal degradation [23, 27, 28]. This inhibition results in the stabilization of HIFs, allowing them to accumulate and activate their downstream effects. HIF stabilization then increases the expression of hypoxia response element (HRE) genes, which creates a cytoprotective environment and decreases cell death in hypoxic conditions. However, it remains unclear how DFO affects SC protein expression, phenotype, morphology, survivability, and transcriptome in response to ROS-induced cell death. While it is ascertained that DFO provides protection against ferroptosis [29, 30], questions remain on whether it is effective in preventing acute cell death mediated by ROS.

The efficacy of DFO as a protective agent against oxidative stress has previously been demonstrated in serum-containing conditions, where it has shown potential to stabilize the expression of hypoxia inducible factor (HIF) family proteins and protect cells under hypoxic conditions, thereby reducing ROS-induced cell death [23, 27, 31, 32]. However, the relevance of serum-based treatments to modern clinical practices is limited, due to the inherent variability in the composition of serum [33–36]. A serum-free approach with predefined and consistent composition offers a more reliable alternative, enabling reproducible and clinically relevant results. Hence, it is important to evaluate the effects of DFO on SCs in serum-free condition to determine whether its protective properties remain effective and clinically relevant.

Here, we investigated the effectiveness of DFO in increasing the survival of Schwann cells (SCs) upon hydrogen peroxide-induced cell death in an in-vitro environment designed to simulate clinical transplant conditions. We hypothesized that SCs pre-treated with DFO would exhibit increased survival in ROS-induced cell death in serum-containing conditions. Additionally, we hypothesized that the neuroprotective effects of DFO would persist in serum-free conditions, such as chemically defined media (CDM), which eliminates the need for patient serum extraction while decreasing serum facilitated SC variation in vivo. We examined the efficacy of DFO in both serum-containing (CDM) and serum-free (CDM) conditions (supp. Table 1), and assessed our results through phenotypic staining, cell count, whole cell corrected total cell fluorescence analysis and RT-qPCR cell death transcriptomics. Our findings show a significant increase in cellular survival upon hydrogen peroxide challenge at 62.5μM in populations pre-treated with DFO only in serum-containing conditions. Furthermore, we observed that serum free media (CDM) exhibited inherent neuroprotective properties, presenting it as a viable alternative to serum-containing media altogether.

## 2. Materials and Methods

### 2.1 Generation and Expansion of Schwann Cell Culture

Schwann cells (SCs) were isolated from the ventral roots of 6-month-old adult GFP transgenic Sprague Dawley rats according to our established protocols for SC culturing from immediately dissociated nerve tissues [37]. The initial cell harvest was plated in the form of droplets directly onto PLL-laminin-coated dishes and expanded in DMEM medium supplemented with 10% FBS, heregulin (10 nM), and forskolin (2 uM) up until confluency. Cells were subsequently lifted from their dishes by trypsinization and purified of contaminating fibroblasts by MACS sorting after incubation with Thy1.1. antibodies, as described in Ravelo et al. [38]. The purified SCs were expanded up to passage-1 (P1) in the abovementioned medium for the creation of cryogenic stocks [37]. These stocks were maintained in liquid nitrogen until use. Prior to experiment, P1 SCs were expanded in D10S 3F (supp Table 1). During expansion, SCs were cultured in poly-L-lysine (PLL, Millipore Sigma P6282) coated 100 mm Corning-treated tissue culture dishes and fed with 7 mL of CDM 3F every 2-3 days. Once the desired confluency (approx. 5 million cells per plate) and maturation state was achieved by visual confirmation of cell swirling, SCs were either passaged into new 100 mm Corning-treated tissue culture dishes or passaged into experimental conditions. SCs were detached using 0.05% trypsin-EDTA (Gibco 25300054) and seeded into PLL coated four-well chamber slides (Nunc Lab-Tek II Chamber Slide System, ThermoFisher), at 50,000 cells per well, for immunocytochemistry experiments and fed with 350 μL CDM 3F per well every 2-3 days until start of experiment (Section 2.3). For RT-qPCR experiments, SCs were plated into 60 mm PLL-coated TPP tissue culture dishes (TPP 93060) at 450,000 cells/plate and fed with 5 mL of CDM 3F every 2-3 days until treatment (Section 2.4). All SCs were cultured in a 5% CO2, 37oC incubator except for the cells at passage-zero (P0) and P1, which were cultured in a CO2 incubator set up at 9% CO2. P2-P5 SCs were used for all experiments. The purity of the SC populations was >98% at P1, as judged by co-immunostaining with S100B and Thy1.1 antibodies. SCs from ventral roots were highly proliferative and visually indistinguishable from the ones derived from sciatic nerve [37].

### 2.2 Experimental treatments and timeline

SCs were plated onto four-well chamber slides for phenotypic staining and subsequent onto 60 mm PLL-coated TPP tissue culture dishes for cell death transcriptomics analysis. All SCs are initially cultured in serum containing CDM 3F media until the desired confluency was reached by visual confirmation of swirling. Prior to 62.5 μM H2O2 challenge or sham, SCs were treated with their respective pre-treatment conditions (supp Table 2) for 24 hours, followed by a 16-hour exposure to 62.5 μM H2O2 in base media. 3F signaling was cleared for selected groups (Fig 3, Fig 4) comparing SCs serum and serum-free. SCs were fed with their respective base medium for 4 days, figures 3A & 4A illustrates medium changes. For sham, SCs were refed with base media only. After the treatment period, SCs were either fixed with 4% paraformaldehyde for subsequent phenotypic staining and density analysis or harvested as cell pellets for RT-qPCR cell death transcriptomics analysis (Section 2.4). Figure 1A illustrates the experimental timeline and figures 1B, 1C shows the effect of the 16-hour exposure to 62.5 μM H2O2 on SCs. Figures 2A, 3A, 4A, 7A show the timeline of the media condition changes during the experiments.

**Figure 1.**
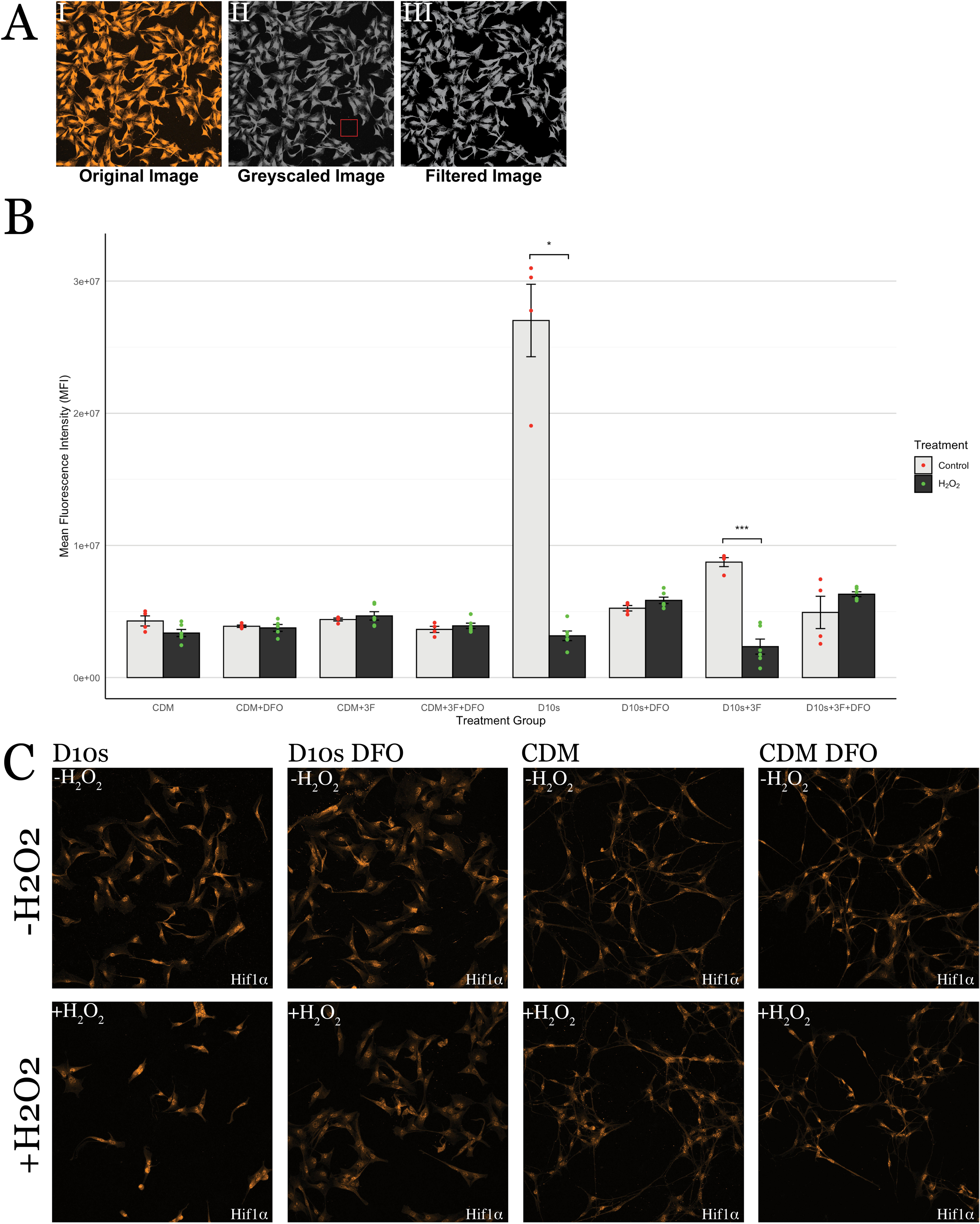
Schematic of experimental design and Schwann Cell Cultures. **(A)** Schematic of the experiments. Timepoints of treatments and hydrogen peroxide challenge. **(B)** Representative brightfield images of Schwann cells 16 hours following hydrogen peroxide challenge **(Biii, iv)** or sham **(Bi, ii)** in serum containing medium. **(Bi, iii)** shows 20x image and **(Bii, iv)** shows 40x image. **(C)** Representative brightfield images of Schwann cells 16 hours following hydrogen peroxide challenge **(Ciii, iv)** or sham **(Ci, ii)** in serum free medium. **(Ci, iii)** shows 20x image and **(Cii, iv)** shows 40x image. All images are taken on Invitrogen EVOS XL CORE inverted brightfield microscope using EAchro 40x LWD PH, 0.65NA/3.1WD (AMEP4635) and Achro 20 x LWD PH, 0.40 NA/6.8 WD (AMEP4934) objectives.

**Figure 2.**
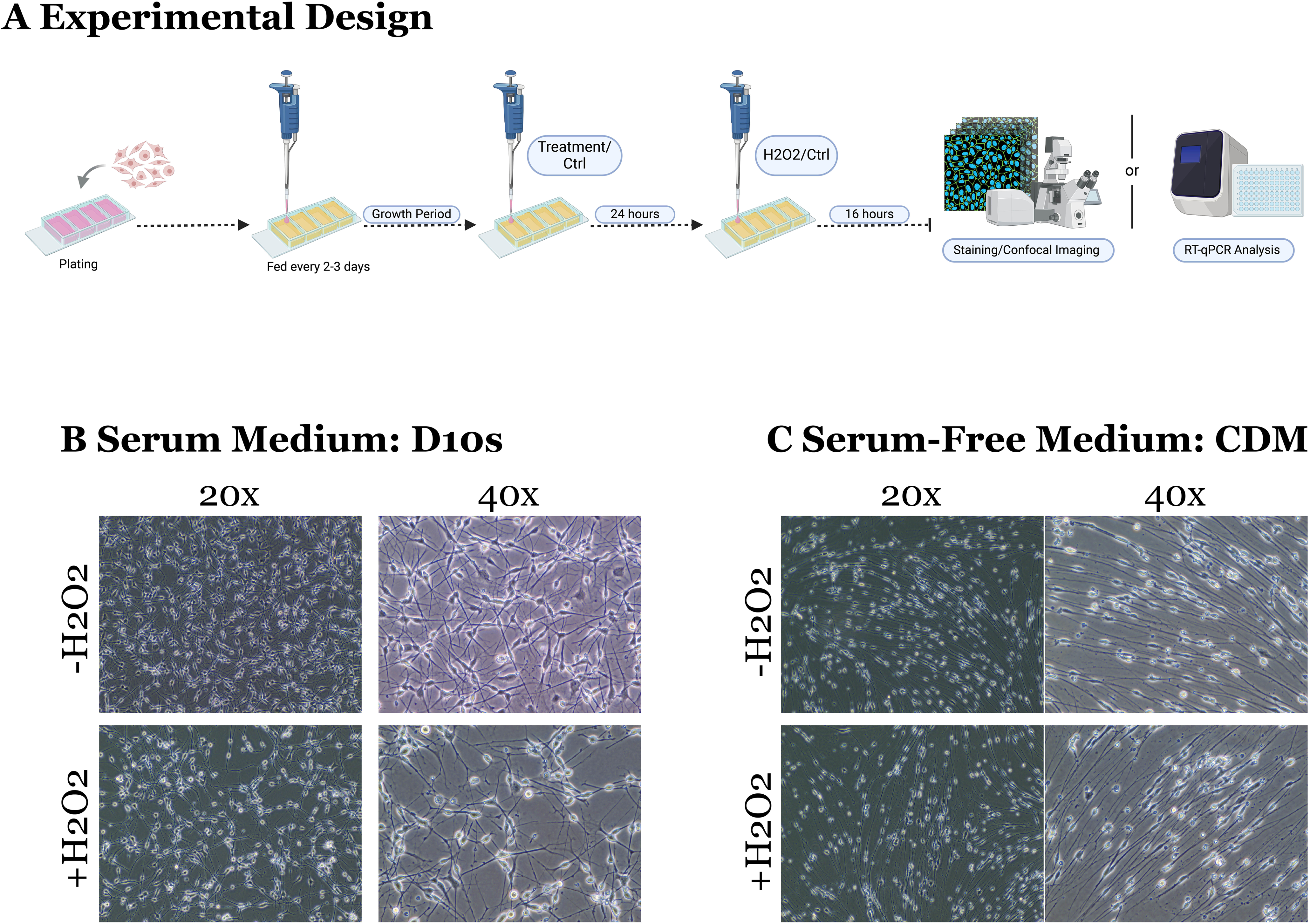
Neurotrophins and growth factor do not increase Schwann Cell survival. **(A)** Schematic of the experiment. SCs were grown for 4 days on 4 well chambers prior to exposure to treatment conditions. **(B)** Quantification of cellular survival in sham versus hydrogen peroxide challenged SCs in D10s Mit, D10s 3F, D10s Mit +bFGF, D10s Mit +FGF5 shows no effect of neurotrophins and growth factors in increasing SC survival. **(C)** Representative images of sham SCs in different conditions. Asterisk labels fibroblast within SC cultures. Images are taken at x40. Channels: DAPI (405, blue), GFAP (488, green), P75 (Cy3, orange). Scale bar = 50um.

**Figure 3.**
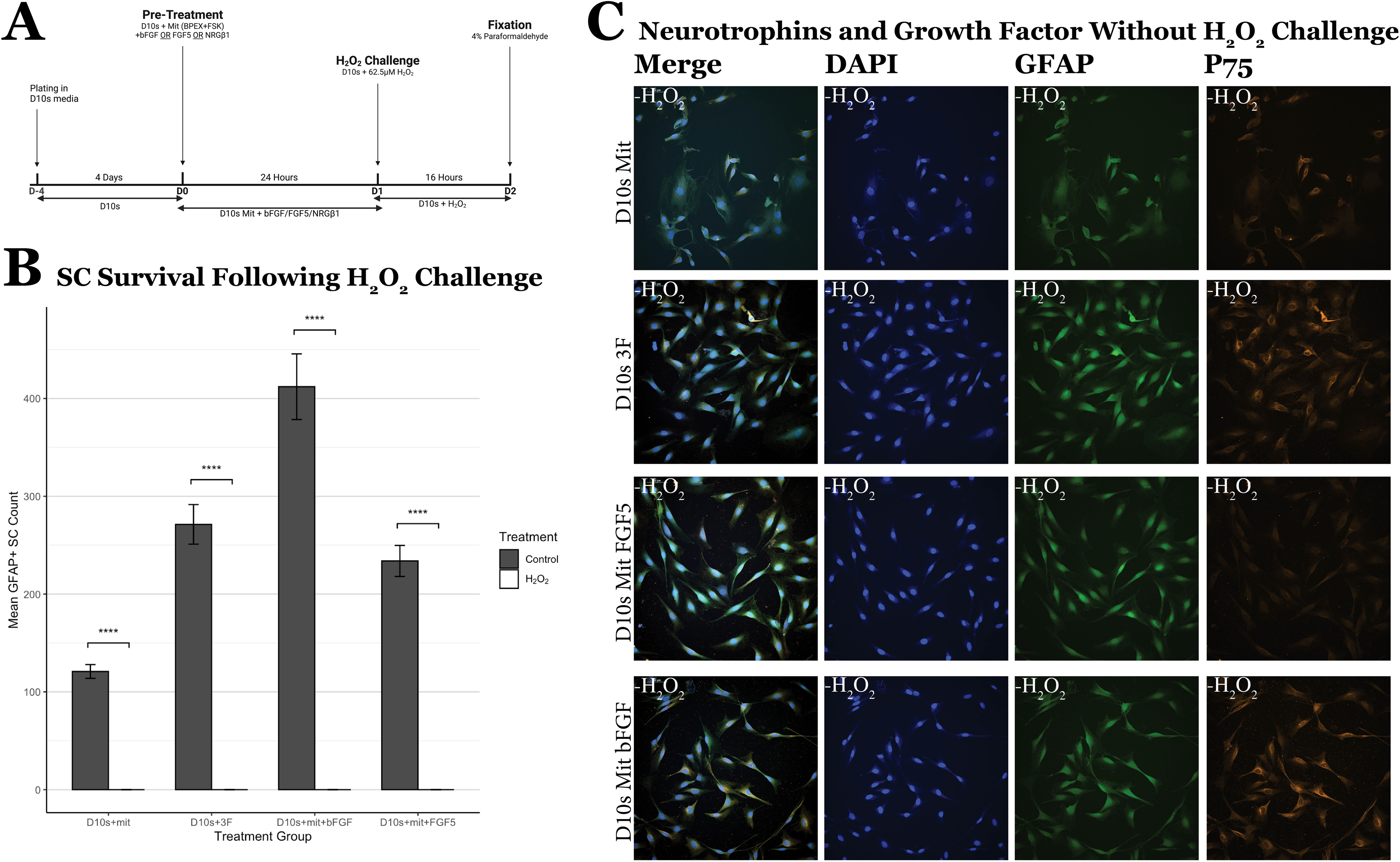
DFO increases Schwann Cell survival in Serum Containing Medium. **(A)** Schematic of the experiment. SCs were grown for 8 days in D10s 3F medium to eliminate fibroblast in culture and promote SC survival and proliferation. **(B)** Quantification of GFAP+ SC nucleus 16 hours following sham or hydrogen peroxide challenge. D10s +DFO survival significantly higher compared to D10s. D10s 3F +DFO survival significantly higher than D10s 3F. **(C)** Representative images of D10s and D10s +DFO following 16 hours of 62.5uM hydrogen peroxide challenge. Channels: DAPI (405/blue), GFAP (488, green), Hif1a (cy3, orange). **(D)** Representative images of D10s and D10s +DFO following 16 hours of sham. Channels: DAPI (405/blue), GFAP (488, green), Hif1a (cy3, orange). Hif1a stain shown in fig 5. Images taken at 20x magnification and 2x zoom.

**Figure 4.**
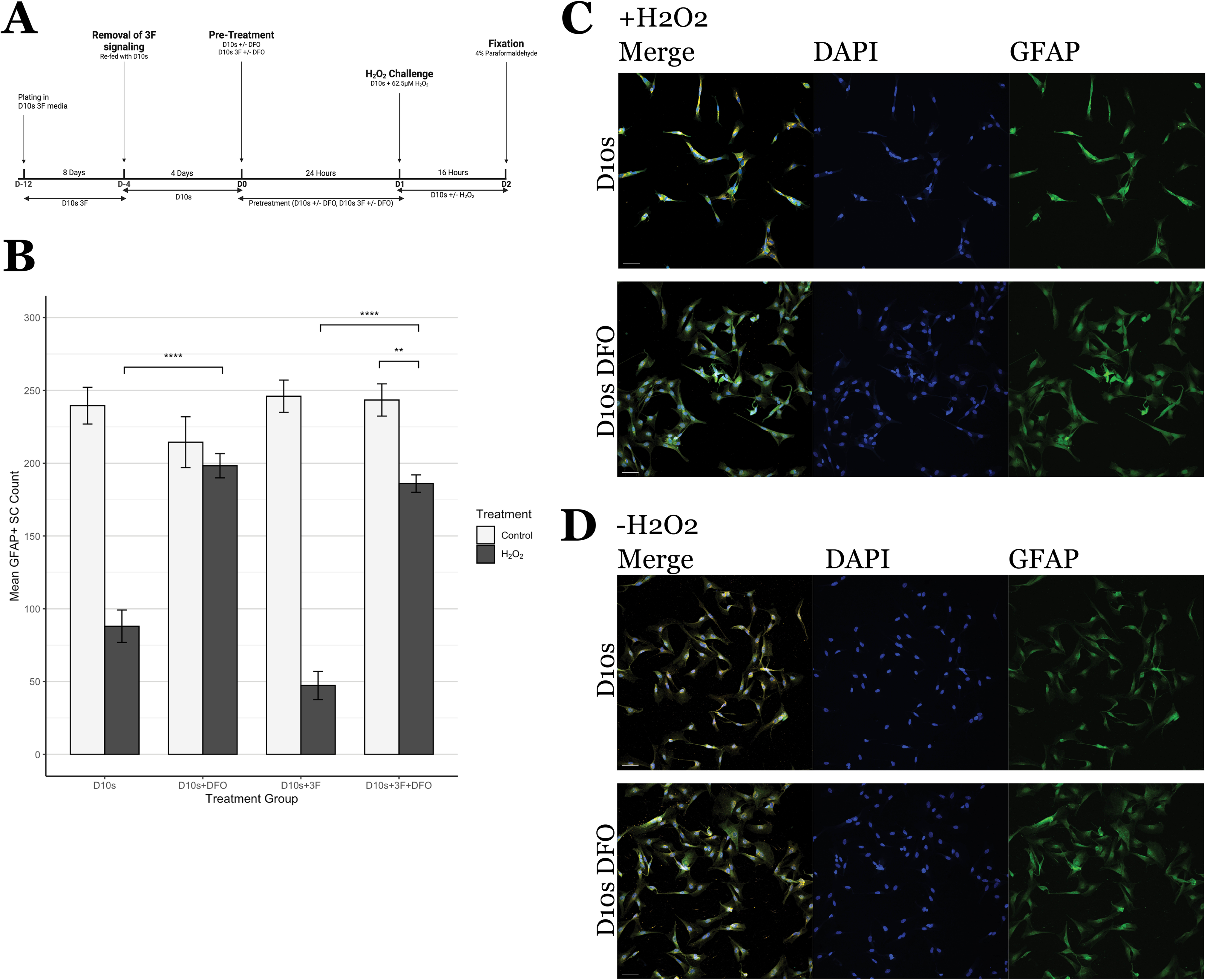
Serum free medium protects Schwann Cells in H2O2 induced cell death. **(A)** Schematic of the experiment. SCs were grown for 8 days in D10s 3F medium to eliminate fibroblast in culture and promote SC survival and proliferation. **(B)** Quantification of GFAP+ SC nucleus 16 hours following sham or hydrogen peroxide challenge. CDM +DFO survival shows no significant increase compared to CDM. CDM 3F +DFO survival shows no significant compared to CDM 3F. **(C)** Representative images of CDM and CDM +DFO following 16 hours of 62.5uM hydrogen peroxide challenge. Channels: DAPI (405/blue), GFAP (488, green), Hif1a (cy3, orange). **(D)** Representative images of CDM 3F and CDM 3F +DFO following 16 hours of sham. Channels: DAPI (405,blue), GFAP (488, green), Hif1a (cy3, orange). Hif1a stain shown in fig 5. Images taken at 20x magnification and 2x zoom.

**Figure 5.**
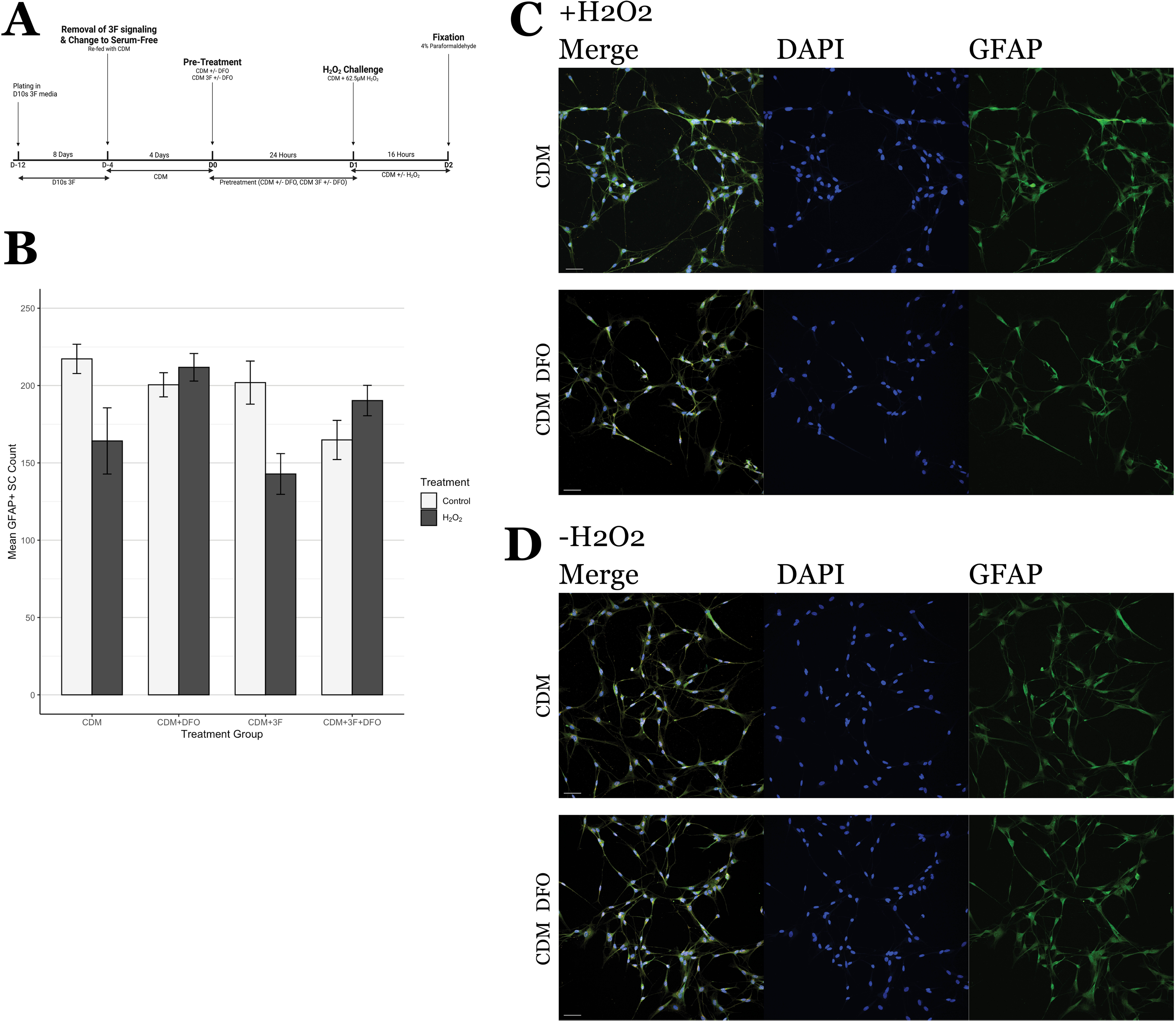
Hif1a WC-CTCF reveals serum variability. **(A)** Representative process of fluorescence intensity quantification. **(Ai)** Original image without processing. **(Aii)** Greyscale image, box shows definition of “background”. **(Aiii)** Image after background filtering to eliminate noise. **(B)** Quantification of Hif1a expression across groups. Statistics shows high variability in serum groups with significant upregulation of Hif1a in non-DFO pretreatment groups. **(C)** Representative images of Hif1a expression across groups.

### 2.3 Immunofluorescence Staining

16 hours after H2O2 challenge, SCs were fixed via 15 minutes incubation with approximately 300 μL of chilled 4% paraformaldehyde in room temperature. After PFA incubation, SCs were washed three times with 1X PB and stored in +4 oC with phosphate buffer (PB) until immuno-staining. We stained SCs with GFAP (Chicken, 1:1000, Aves GFAP), P75 (Rabbit, 1:400, Bioss BS-0161R), HiF1α (Rabbit, 1:500, Sigma Aldrich SAB5701087), and Collagen IV (Rabbit, 1:400, Rockland 600-401-106-0.1) primary antibodies. Secondary antibodies used were from Jackson Immuno and used at 1:800, we used donkey anti-chicken AlexaFlour 488 (703-546-155) and donkey anti-rabbit cy3 (711-166-152). SCs were incubated at room temperature with primary antibody in blocking solution: 90% 1X phosphate buffer (PB), 10% Donkey Serum (Lampire 7332100), and 0.2% Triton X-100 (Sigma Aldrich X100), for 2 hours, then washed with 1X PB three times. SCs are then incubated with secondary antibody in blocking solution at room temperature for 45 minutes in the dark. Cells are subsequently washed with 1X PB three times and cover slipped with Prolong Gold or Diamond Antifade Mountant with DAPI (Invitrogen).

### 2.4 Confocal microscopy

All imaging was performed on a Nikon AXR confocal microscope using the Nikon NIS Elements software. The microscope is equipped with four lasers, ranging from 405-647. All cells were stained with DAPI (405) and their respective stains for qualitative and quantitative analysis (GFAP, P75, HIF1α, Collagen IV). Image acquisition parameters are described in the sections below, in the context of their application. Representative images taken for figure 2 was taken using Nikon APO LWD 40XxWI lS DIC N2 (NA=1.15, WD=610μm) objective. Representative images for figures 3-6 was taken using Nikon APO LWD 20x WI lS (NA=0.95, WD=950μm).

### 2.5 Survival Quantification

Images taken from the wells were quantified using the NIH ImageJ software. Quantification was semi-automated with human correction of software errors. Images were taken at 10x (NA=0.45, WD=4000μm) magnification for each well. For each well, we randomly selected 3-4 fields. Using image J, ND2 images were opened as gray-scale images. The detection threshold was manually adjusted for all images to accurately highlight each individual DAPI stain. Images were then converted to binary images to remove background noise before being processed with the watershed algorithm to split any combined nucleus. DAPI nuclei were subsequently quantified using particle analysis with the size detection threshold set to 100 pixels to infinity. Quantification results were then compared with the original DAPI and GFAP stain manually to ensure accurate quantification and address any potential errors (eg: splitting of a single nucleus, grouping of multiple nuclei, counting GFAP negative nuclei). Final quantifications were manually corrected for any software errors.

### 2.6 Hif1a & Collagen IV Quantification

Immunofluorescence images were obtained via confocal microscopy described in section 2.5. Images were taken using a 20x objective (NA=0.75, WD=1000 μm), maintaining identical parameters for all Hif1a and Collagen IV scans as follows: laser power: 33, gain: 84.2, pinhole size: 1.2, zoom: 1x, z-stack: 50 μm, and step size: 2 μm. For Hif1a, we selected n=6 per pre-treatment group (supp. Table 2). For collagen IV, we selected n=4 per pre-treatment group. The images were then exported as ND2 files and further processed using a custom, semi-automated pipeline to quantify whole cell corrected total cell fluorescence (WC-CTCF) using FIJI/ImageJ software. In brief, ND2 files were split into single channels and compressed to maximum intensity projections (MIPs) for quantification. Then, the MIPs for Hif1a and Collagen IV were measured for total fluorescence in the image (integrated density), subtracted by the total background fluorescence. Utilizing this approach permits correction for size, density, or morphological differences between cells in their respective treatment groups. The formula used was CTCF=Integrated density -(total area × background intensity mean). We analyzed the whole cell fluorescence level to capture cytoplasmic and nuclear expression levels.

### 2.7 Statistical Analysis

All statistical analyses were conducted using the RStudio software (2024). Data were presented with error bars indicating Standard Error of Mean (SEM). Depending on data characteristics, either Welch’s one-way analysis of variance (Welch’s ANOVA) or standard ANOVA was used for F-tests. Post hoc comparisons were performed using Tukey’s HSD, Dunn’s Test, or Games-Howell test, as appropriate. p ≤ 0.05 denotes statistical significance. Significance was also presented with the following notations: * (p ≤0.05), ** (p ≤0.01), *** (p ≤0.001), **** (p ≤0.0001). For RT-qPCR analysis, fold change was reported on the log2 scale, and comparative analyses were performed on GeneGlobe (Qiagen).

### 2.8 Cell death transcriptomics 2.8.1 RNA Isolation

Selected groups (Supp. Table 3) were chosen for transcriptomic analysis. Samples were lysed using the Qiashredder (Qiagen 79656). RNA was isolated using RNeasy Micro Kit (Qiagen 74004) following manufacturer’s instruction. Isolated RNA was subsequently stored in –80 oC until use. The quantity of extracted RNA was evaluated using Nanodrop 2000 (ThermoFisher) or BioTek Synergy HTX Multimode Reader (Agilent), while the integrity was evaluated using Bioanalyzer 2100 (Agilent) for RNA Nano Chips (Agilent) and TapeStation 4200 (Agilent) for RNA ScreenTapes (Agilent). RNA integrities were ensured to be of sufficient quality prior to PCR. 100 ng of total RNA was utilized for subsequent analysis within each group.

#### 2.8.2 cDNA synthesis

cDNA was synthesized using RT2 First Strand Kit (Qiagen, 330404) following manufacturer’s instructions. Samples were processed using the T100 Thermo-Cycler (BioRad) starting with a 15-minute incubation at 42 °C followed by a 5 minute denaturation at 95 °C. Synthesized cDNA was stored at –20 oC until use. The quality and quantity of synthesized strands were evaluated using Nanodrop 2000 (ThermoFisher).

#### 2.8.3 qPCR

The 96-well RT2 Profiler PCR Array Rat Cell Death PathwayFinder (Qiagen 330231 PARN-212ZA) was used to detect expression levels of 84 targeting genes and 5 control genes (supp. table 4). Selected genes were essential for the central mechanisms of cellular death, spanning apoptosis, autophagy, and necrosis. Following manufacturer’s instructions, cDNA was mixed with RT2 SYBR Green qPCR Mastermix (Qiagen 330502) before loading onto the sample plate. The sample plate was processed using the QuantStudio3 System (Applied Biosystems, Inc.) starting with a 10-minute incubation at 95 °C, followed by 40 cycles of denaturation at 95 °C for 15 seconds and annealing/extension at 60 °C for 30 seconds. Geneglobe (Qiagen) was used for comparison studies, fold change regulation and heatmap generation. The detection threshold was standardized to 20,000. The CT threshold is set to 35 and control genes were normalized using geometric means.

#### 2.8.5 Gene Ontology Analysis

A gene ontology (GO) analysis was performed based on the heatmaps generated from RT-qPCR data. Differentially expressed genes were quantified using log2fold change and reported as up or downregulated, relative to a CDM control. Groups compared CDM vs CDM, both without H2O2 and CDM vs D10S, both with H2O2 and DFO treatment. GO enrichment analysis was performed using PANTHER Classification System (v19.0, pantherdb.org). The individual genes were entered into the database, specifying rattus norvegicus as the reference species. Functional annotation categories included biological process, molecular function, cellular component, and protein classification. Due to biological replicates of n=1 per condition, statistical testing for the RT-qPCR results was not conducted.

## 3. Results

### 3.1 Variation in SC Morphology between Serum-Containing and Serum-Free Conditions

Schwann cell morphology varied between serum-containing and serum-free conditions, with distinct differences observed in soma and cellular processes. In serum-containing conditions, SCs exhibited a more expanded soma, which appeared flatter and covered a larger surface area (Fig. 1B). In contrast, SCs cultured in serum-free conditions displayed a more compact, needle-like bipolar soma characterized by a narrower and more defined structure (Fig. 1C). Similarly, differences were observed in the morphology of SC processes. In serum-containing conditions, SCs processes were comparatively shorter and remained proximal to the soma with limited extension (Fig. Fig. 1B). In contrast, SCs in serum-free conditions displayed more extended, fiber-like processes that stretched distally and forms bipolar processes from the soma, indicative of a more differentiated state (Fig. 1C). SCs in serum-containing and serum-free conditions also responded differently in the event of H2O2-induced oxidative stress. In serum-containing conditions, SC morphology and density were greatly altered in the presence of H2O2-challenge (Fig. 1B), whereas a more consistent SC phenotype was observed in serum-free conditions (Fig. 1C)

### 3.2 Growth Factors Failed to Promote SC Survival Following H2O2 Exposure

We investigated the potential of growth factors to enhance cellular survival following H2O2-induced oxidative stress. By following the experimental paradigm (Fig. 2A), SCs were pre-treated with either basic fibroblast growth factor (bFGF), fibroblast growth factor 5 (FGF5), or Neuregulin Beta1 (NRG) - in D10s Mit media (supp. Table 2) for 24 hours prior to H2O2 exposure. Following their pretreatments, SCs were analyzed by immunofluorescence staining to evaluate DAPI, p75, and GFAP expression (Fig 2C). SC survival was evaluated by quantifying GFAP-positive SC nuclei, followed by statistical analysis. Comparing growth factors pretreated SCs without H2O2 control, D10s Mit, shows significantly increased proliferation (Fig 2B; D10s + Mit vs D10s + Mit + bFGF, p= 7.26 x 10-22, ****; D10s + Mit vs D10s + Mit + FGF5, p= 3.19 x 10-4, ***; D10s + Mit vs D10s + 3F, p= 4.76 x 10-7, ****), validating their role in promoting proliferation of SCs. However, SCs were not protected against H2O2 induced cell death when pre-treated with neurotrophin or growth factors, as differences in GFAP-positive SC nuclei across treatments were statistically insignificant (Fig 2B), indicating that pretreatment with growth factors are not sufficient to protect SCs against reactive oxygen species induced cell death (D10s + Mit control vs D10s + Mit + H2O2, p= 1.51 x 10-2, *; D10s + Mit + bFGF control vs D10s + Mit + bFGF + H2O2, p= 7.26 x 10-22, ****; D10s + Mit + FGF5 control vs D10s + Mit + FGF5 + H2O2, p= 1.63 x 10-9, ****; D10s + 3F control vs D10s + 3F + H2O2, p= 1.09 x 10-10, ****). Morphological observations from confocal images of SCs grown in selected growth factors (Fig. 2C) confirmed that growth factors promoted the typical elongated morphology of *in vitro* cultured SCs.

### 3.3 Deferoxamine Mesylate (DFO) Increased SC Survival Following H2O2 Challenge in Serum Containing Conditions

Deferoxamine Mesylate (DFO) is an FDA approved iron chelator previously suggested to enhance Schwann cell survival in hypoxic conditions [22]. We tested if DFO could effectively prevent H2O2-induced SC death in serum-containing conditions (D10s). We also tested whether the addition of 3F during DFO treatment would influence DFO’s protective effect against H2O2-induced oxidative stress. We hypothesized that 3F would prime SCs for growth and proliferation, which could, in turn, impair their ability to response to H2O2 induced cell death. Following their treatment protocols (Fig 3A, 4A), SC survival was evaluated by quantifying GFAP-positive SC nuclei, followed by statistical analysis via Tukey’s HSD test. Results showed that, under H2O2-induced cell death, SC did not survival in D10s (Fig. 3B; D10s + H2O2 vs D10s -H2O2, p= 1.15 x 10-13, ****). Additionally, inclusion of 3F did not lead to any significant improvement in SC survival (Fig. 3B; D10s + 3F + H2O2 vs D10s + 3F Control, p= 3.73 x 10-14, ****), consistent with our previous findings that 3F does not provide any protective effect against oxidative stress (Fig 2B; D10s 3F +H2O2 vs D10s 3F control, p= 1.09 x 10-10, ****). We added DFO to evaluate its effects on SC survival and phenotype in the presence of H2O2-induced cell death. The inclusion of DFO in both D10s and D10s 3F medium protected SCs and increased survival following exposure to H2O2 (Fig. 3B; D10s + H2O2 vs D10s + DFO + H2O2, p= 2.39 x 10-10, ****; D10s 3F + H2O2 vs D10s 3F + DFO + H2O2, p= 1,18 x 10-13, ****). Furthermore, comparisons between D10s DFO +H2O2 and D10s -H2O2 showed no significant difference in SC survival, indicating that DFO reduced H2O2-induced cell death and restored SC survival to baseline levels. However, in D10s 3F medium, DFO pre-treatment (D10s 3F +DFO +H2O2) did not protect SC survival to baseline levels (D10s -H2O2). This suggests that while 3F is essential in promoting SC proliferation, it can interfere with the effects of DFO and reduce its protectiveness. Nonetheless, these results showed that DFO conferred significant protection against H2O2-induced cell death in serum-containing conditions. Immunofluorescent staining indicated that H2O2-challenge resulted in altered SC morphology and non-uniform SC distribution in serum-containing conditions (Fig. 3C)

### 3.4 Serum-Free Media Protected SCs Against H2O2-Induced Cell Death without DFO

We also investigated if DFO could effectively prevent H2O2-induced cell death in a serum-free, chemically defined media, a DMEM/F12 medium supplemented with bovine insulin, human transferrin, sodium selenite, and putrescine (Supp. table 1). The experimental protocol and quantification methods outlined in section 3.3 were repeated with the only modification being the substitution of serum-containing D10s with serum-free CDM as the base medium (Fig 4A). Interestingly, CDM alone was sufficient to promote SC survival under H2O2 induced cell death (Fig. 4B, CDM -H2O2 vs CDM +H2O2, p=1.157e-13, ns). Even without the addition of DFO, CDM alone restored SC survival to baseline levels, suggesting that factors in CDM may contribute to preventing H2O2-induced cell death. Here, the inclusion of 3F did not decrease the protection provided by CDM against H2O2 induced cell death (Fig 4B, CDM 3F -H2O2 vs CDM 3F +H2O2, p= 5.037e-02, ns), indicating that 3F is not sufficient to interfere with the effects of CDM and reduce its protectiveness, unlike their serum counterpart. The inclusion of DFO in serum-free CDM did not affect SC survival in H2O2 induced cell death, indicating that DFO is unnecessary in serum free CDM to promote SC survival (Fig 4B, CDM DFO -H2O2 vs CDM DFO +H2O2, p= 9.989e-01, ns). Immunofluorescent staining showed SC distribution and morphology was mostly unaffected after H2O2-challenge in serum-free conditions (Fig 4C).

### 3.5 Hif1a expression consistent in serum-free but inconsistent in serum conditions

It is previously reported that DFO acts as a neuroprotective agent through the upregulation of the hypoxia inducible factor (HIF) protein family to promote cellular survival in hypoxic environments such as the ROS rich environment in our study. We tested if Hif1a will be upregulated here with our SCs. To this end, we have stained for hypoxia inducible factor 1 alpha (Hif1a) and subsequently quantified the protein expression using whole cell corrected total cell fluorescence (WC-CTCF). Statistical analysis of the Hif1a WC-CTCF reveals a consistent Hif1a expression across serum-free groups regardless of DFO pretreatment or H2O2 challenge (fig 5B). Hif1a expression in serum shows irregular expression with a significantly high expression of Hif1a in D10s (fig 5B, D10s -H2O2 vs D10s +H2O2, p= 1.800e-02, *) and D10s 3F (fig 5B, p= 2.300e-04, ***) without H2O2 challenge (fig 5B). Qualitative analysis on immunocytochemistry images reveals localized nuclear expression of Hif1a in serum-free conditions and higher levels of cytoplasmic Hif1a expression in non DFO pretreated and non-H2O2 challenged SCs in serum conditions (fig 5C).

### 3.6 Serum Drives High Irregularity in Collagen IV Expression

The secretion profile of the extracellular matrix (ECM) by SCs plays a critical role in SC adhesion, maturation, and subsequent nerve regeneration following transplantation. Collagen IV is one of the key ECM elements expressed by SCs involved in the regeneration of myelin sheath, an indicator of SC maturation [35]. We investigated whether the addition of DFO would influence collagen IV expression profile of SCs under both serum and serum free conditions, with and without H2O2 induced cell death. Collagen IV expression was analyzed through immunofluorescence staining, followed by WC-CTCF analysis to quantify collagen IV expression (section 2.7). WC-CTCF analyses revealed varying patterns of collagen IV expression between SCs in serum-containing and serum-free medium. Collagen IV expression was generally consistent across SCs cultured in CDM. The addition of DFO did not result in any measurable changes in collagen IV expression in majority CDM groups when compared to their non-DFO treated counterparts (supp. table 6), suggesting that DFO did not alter the ECM expression of collagen IV in serum-free conditions. Surprisingly, when SCs are challenged with H2O2 in CDM medium (CDM +H2O2), collagen IV expression shows a significant drop compared to control. Indicating that DFO or 3F may be needed in serum-free conditions to promote the production of extracellular collagen IV in SCs in the presence of ROS. In serum conditions, collagen IV expression is highly variable. When pretreating SCs with 3F, collagen IV expression is highly upregulated without the presence of ROS (Fig 6; D10s 3F -H2O2 vs D10s 3F +H2O2, p= 1.000e-05, ****). However, when pretreatment includes DFO, the upregulation of collagen IV is diminished. Overall, SCs in serum conditions exhibits largely variable collagen IV expression compared to serum-free conditions.

### 3.7 Serum vs serum-free conditions show differential transcriptional responses in cell death pathways

We asked if treatment with DFO leads to transcriptional changes in cell death pathways, specifically apoptosis, necrosis, and autophagy pathways. To this end, we have isolated RNA from SCs in varying conditions (supp. table 8) for use in RT-qPCR to investigate transcriptional changes in cell death pathways. The fold changes of genes were measured as upregulation or downregulation in serum-free (CDM) groups relative to serum-containing (D10S) groups. The largest fold changes when comparing CDM vs D10S without H2O2 were observed in genes associated with necrosis, specifically downregulation of Maco1 (−220.02) and Mag (−24.68). The largest fold change observed in this group was for Olr1583 (20.47), which plays a key role in necrosis. In comparison, DFO and H2O2 treated groups (CDM+DFO+H2O2 vs D10S+DFO+ H2O2) showed the greatest fold changes in both anti-apoptotic genes and necrosis genes. The largest upregulation was seen in genes associated with necrosis, including Defb1 (21.09) and Hspbap1 (11.68). In contrast, the top downregulated genes were Tnfrsf11b (−17.64) and Bcl2L11 (−8.08), both of which are associated with anti-apoptotic pathways. Considering the substantial reduction of Maco1 transcription when comparing CDM vs D10S, genes associated with autophagy, Ins2 (19.87) and Esr1 (17.1), showed the next largest increase in transcription. Overall, the data suggests CDM primarily reduces genes involved in necrosis, while increasing genes associated with autophagy, in the absence of H2O2. When both groups are treated with DFO and H2O2, genes involved in necrosis are upregulated, while anti-apoptotic transcripts are suppressed. Interestingly, fold changes for autophagy genes appear largely unchanged. Heatmaps of RT-qPCR are shown in figures 7 and 8.

**Figure 6.**
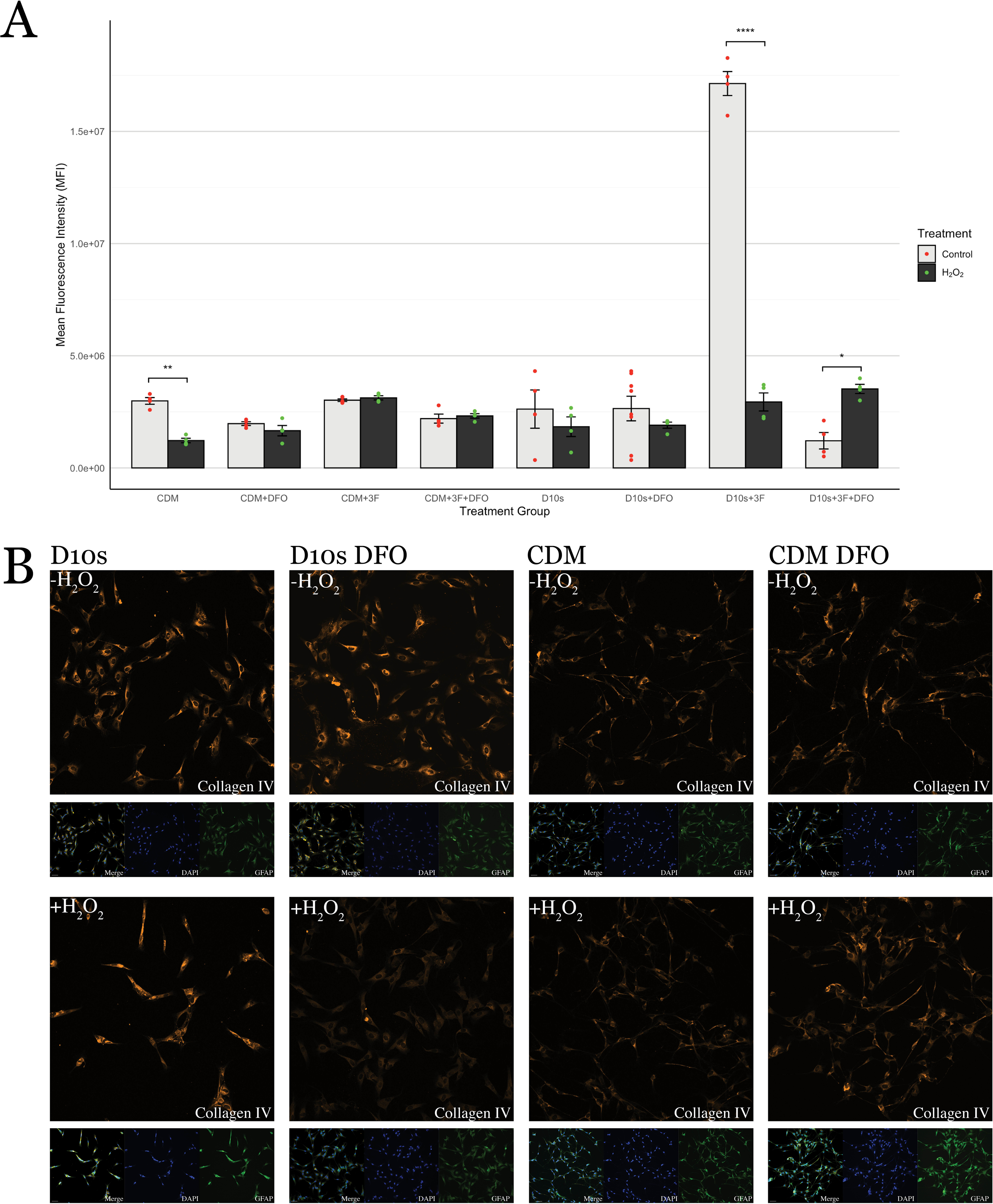
Collagen IV WC-CTCF shows Schwann Cell states. **(A)** Quantification of collagen IV expression. P values for statistics shown in supplemental table 7. **(B)** Representative images of collagen IV expression across groups following 16hours of 62.5uM hydrogen peroxide challenge or sham. Channels: Collagen IV (Cy3, orange), GFAP (488, green), DAPI (405, blue).

**Figure 7.**
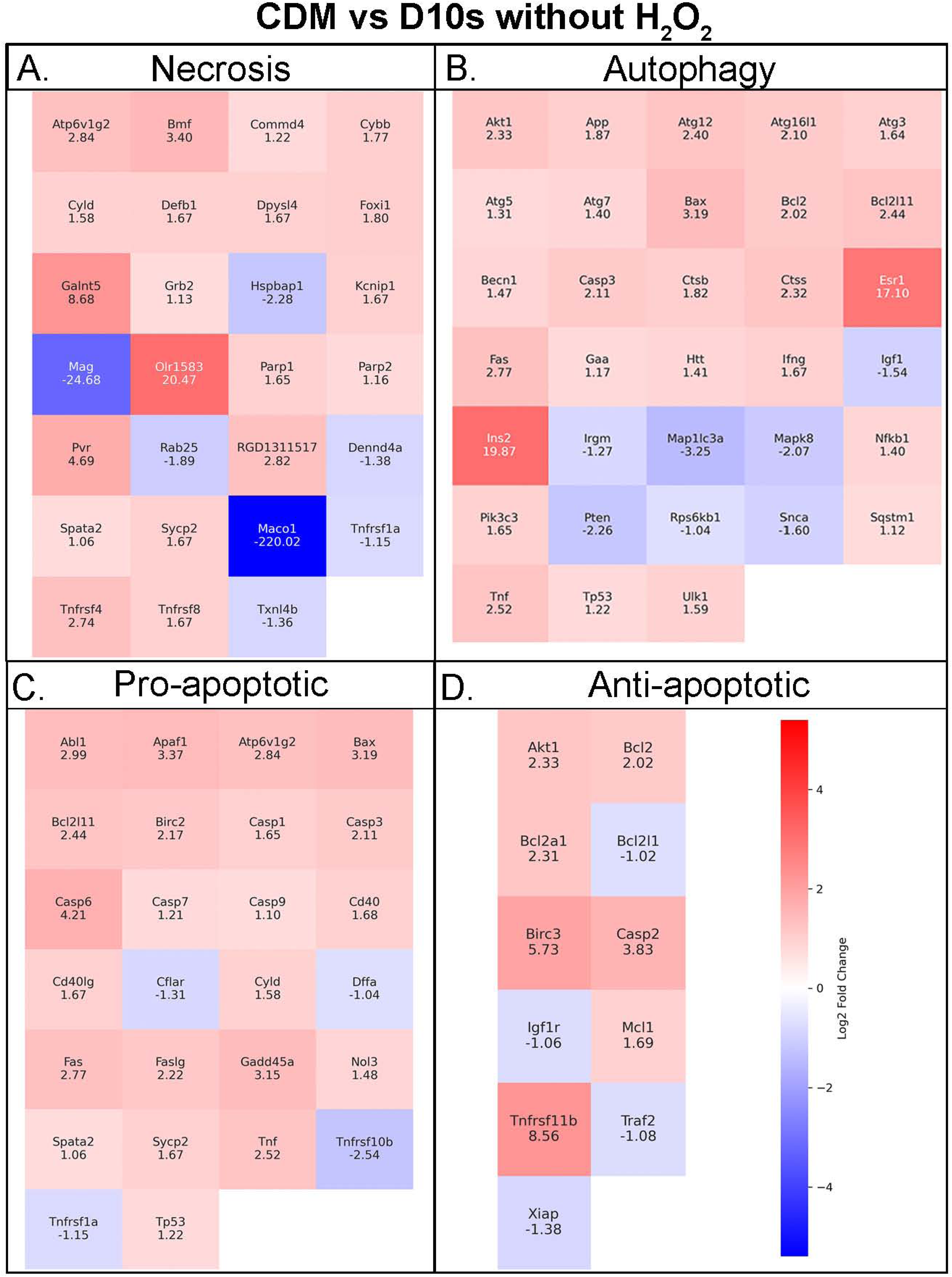
RT-qPCR shows increased metabolism and autophagy in CDM medium. Heatmaps showing Schwann Cell, in serum-free versus serum medium without the presence of H2O2, transcripts expression associated in **(A)** necrosis, **(B)** autophagy, **(C)** pro-apoptotic, and **(D)** antiapoptotic pathways. Fold-change values shown on heatmap.

## 4. Discussion

We examined the effect of Deferoxamine Mesylate (DFO) on the survival of ventral root Schwann cells isolated from adult Sprague-Dawley rats in H2O2-induced cell death. It was previously reported that DFO increases SC survival in H2O2 induced cell death and hypoxic conditions in vitro in serum-containing conditions [23, 42]; here, we expanded the horizon to study DFO’s effectiveness in protecting SCs from H2O2 induced cell death in more clinically relevant, serum-free conditions and presented our results through phenotypic staining, SC survival quantification, whole cell corrected total cell fluorescence (WC-CTCF) analysis, and RT-qPCR cell death transcriptomics analysis.

### 4.1 Distinctive SC Morphology in Serum and Serum-Free

Consistent with previous findings from Plant and colleagues [30, 40], SCs exhibited drastically different morphology between serum-containing and serum-free conditions (Fig. 1). The differential morphology of the two culture conditions affects SC maturation and regenerative capacity in vivo. For instance, the bipolar and spindle morphology of SCs in CDM shows a pro-regenerative phenotype in comparison to the small and flat phenotype of SCs in D10s. The spindle morphology allows for neurites to grown on top, acting as a bridging mechanism and allowing for nerve regeneration. However, SCs in serum conditions switch from a regenerative state into a proliferative state and abandons cell contact [41]. Thus, SCs grown in serum appear to be more prone to H2O2 induced cell death due to the energetic switch to a proliferative state. Furthermore, serum has been reported to lead to unstable and variable phenotypes in vitro in mesenchymal stromal cells and olfactory ensheathing glia [42–44]. Although literature lacks work describing SCs in serum versus serum free medium, the use of serum would still largely decrease the reproductive capacity of SC cultures in transplantation studies due to the lot-to-lot variability in serum production.

### 4.2 Growth Factors Provides no Protection against H2O2 induced cell death

We asked if growth factors would be able to attenuate H2O2 induced cell death in Schwann cells (SCs). We pre-treated SCs with basic fibroblast growth factor (bFGF), or fibroblast growth factor 5 (FGF5), or neuregulin beta 1 (NRGb1), for 24 hours prior to 16 hours of 62.5mM H2O2 challenge. Our results indicate no survival of SCs following H2O2 challenge regardless of pretreatment. Although it has been previously reported that these factors attenuate H2O2 induced cell death in alternative cell types such as embryonic cortical neurons [45] and bone marrow mesenchymal stem cells [46], we have shown that they do not attenuate H2O2 cell death in adult SCs. Here, we did not test these growth factors independently in serum-free conditions. In serum-free growth factor could produce a stronger protectiveness in addition to the serum-free protectiveness through the promotion of proliferation. However, as seen with SCs grown in 3F, it may also drive SCs into growth states and thus reduce their capacity to withstand H2O2 induced cell death.

### 4.3 DFO Efficacy in Serum and Serum-Free Conditions

Here, we have shown that serum-free CDM medium seems to protect SCs from H2O2 induced cell death regardless of DFO pre-treatment. In serum conditions, Hill and colleagues [22, 39] have previously established DFO effectiveness in vitro showing its ability to upregulate hypoxia inducible factor 1 alpha and increased cellular survival, by the means of live dead staining [22]. However, we have shown here that the use of CDM without DFO pretreatment was able to achieve similar levels of protection against H2O2 cell death to DFO in serum-containing D10s medium. The use of CDM is also shown to be a better alternative due to its consistency that is largely unachievable in serum-based treatments due to high variation in serum which contains variable and trace amounts of neurotrophins and growth factors such as BDNF, FGF5, bFGF, and NRG beta 1, which varies from batch to batch in serum production [47]. While DFO has been proven to increase SC survival in serum-containing conditions [22], most of its effects are masked by the inherent protectiveness provided in CDM, suggesting that the inclusion of DFO in CDM may not be necessary to achieve H2O2 induced cell death protection.

### 4.4 Transferrin in CDM medium may influence HiF-PhD to override DFO effects in SCs

The combined factors transferrin, insulin, and sodium selenite, also known as ITS serum free supplement, is a long-established serum free alternative to the use of fetal bovine serum supplement in tissue culture medium. These factors are critical in supporting cellular proliferation, survival, and glucose uptake. Specifically, transferrin acts as an iron transporter and has been reported to decrease iron accumulation in Parkinson’s disease affected neurons [48]. Mechanistically, transferrin receptor (TfR) acts in downstream of Hif1a expression, allowing cells to uptake higher levels of iron during low oxygen environments for increased mitochondrial activity [49, 50]. With the addition of transferrin in the serum-free medium, the process of Hif1a modulated upregulation of TfR may be overwritten by the presence of transferrin to sufficiently provide iron to SCs and may shut down DFO facilitated HiF-PhD inhibition with increased iron levels leading to our observed levels of SC Hif1a expression in serum-free conditions (fig 5C). It may be of interest to pursue the individual effects of each supplement component in H_2_O_2_ induced SC death to further characterize the protection offered in CDM medium.

### 4.5 Nuclear expression of Hif1a facilitate protection in serum-free cultured or DFO pre-treated SCs

We have quantitated whole cell Hif1a expression via WC-CTCF. Here we see a consistent nuclear expression of Hif1a in SCs grown in serum-free conditions (fig 5B, C) but with an inconsistent expression of Hif1a in SCs grown in serum conditions (fig 5B). Furthermore, ICC reveals a cytoplasmic expression of Hif1a in non H2O2 challenged SCs in serum conditions (D10s -H2O2 and D10s 3F -H2O2, Fig 5C). Previous work has reported the expression of Hif1a enhances SC survival when transplanted into the spinal cord [48]. Stabilization and subsequent nuclear translocation of Hif1a allows for binding with hypoxia response element (HRE) genes and induces protection against oxidative stress through the upregulation of critical angiogenic and protective genes such as vascular endothelial growth factor, or VEGF [51, 52, 53], which has previously been reported to be upregulated through western blot and qPCR analysis [22]. Here, our data adds strong evidence for this protection such that the DFO pre-treated SCs survived compared to non-DFO pretreated SCs in serum conditions. However, in serum free conditions, this protectiveness seems to be mediated by an alternative mechanism since regardless of DFO pretreatment or not, SCs survived the H2O2 challenge. Further, the nuclear expression of Hif1a seems to be present in SCs in serum-free conditions but further enhanced by the presence of DFO. It may be crucial to determine the mechanism of this protection to understand the protection of SCs against H2O2 induced cell death in serum-free conditions. It would be of interest to explore the differences of Hif1a presence in the cytoplasm and nucleus in serum versus serum-free conditions as well.

### 4.6 Collagen IV expression reveals proliferative SCs in serum media and pro-regenerative SCs in serum-free media

We quantitated whole cell fluorescence of collagen IV in SCs to determine if there are any extracellular matrix changes. Collagen IV is an important factor in determining states of SCs, it serves as an indicator of proliferation and remyelination potential of SCs. We have shown here that by inducing proliferative states in SCs via the addition of the 3F (forskolin, bovine pituitary extract, and neuregulin) in serum containing medium, survival in H2O2 induced cell death is negatively affected such that survival is decreased compared to SCs without proliferative 3F signaling in serum conditions (Fig 4B, D10s 3F -H2O2 vs D10s 3F +H2O2, p=3.730e-14). Subsequent collagen IV quantitation also reveals a higher proliferative state in D10s+3F medium condition through a high collagen IV expression when not challenged with H2O2, where the proliferative state is shut down following H2O2 challenge (Fig. 6, D10S 3F -H2O2 vs D10S 3F+H2O2, ****, p=1.000e-05). Previous work has shown immature or proliferative Schwann cells do not facilitate regeneration as efficient as matured Schwann cells [26], our results thus indicate that the more mature states of Schwann cells in serum-free media will facilitate higher regeneration as a cellular transplant, owing to the more stabilized extracellular matrix (fig. 6) and their bipolar process phenotype, which allows for host neurons for bridging the lesion gap (fig 2, 4).

### 4.7 Serum free, chemically defined media shifts SCs towards heightened metabolism and cellular recycling phenotype without the presence of H2O2 and DFO

Here, we identified numerous differentially expressed genes in our in vitro model of hydrogen peroxide-induced cell death. Transcriptomic changes associated with necrosis, autophagy, and apoptosis were observed and the GO enrichment analysis has provided key insights into how these genes affect cell death pathways. In the absence of DFO or H2O2, the serum-free group, CDM -H2O2, primarily showed changes in genes of the necrotic pathway, especially downregulation of Maco1 and Mag. In addition to the necrosis pathway, these genes play a role in cytoskeletal dynamics, cell adhesion, myelination, and metabolic shifts in response to environmental cues [48]. Given their combined and robust downregulation, these data suggest that the SCs may be undergoing a metabolic shift, allocating cellular resources for energy production and autophagy rather than myelin maintenance or proliferation [54, 55]. Within the same treatment group, OLR1583, Ins2, and Esr1 were strongly upregulated, which are involved in the necrosis and autophagy pathways [55]. Although, supplementary roles of these genes support the inference that the SCs are indeed shifting toward a pro-survival phenotype. Aside from smell, olfactory receptors can aid in modulating intracellular signaling in response to a dynamic extracellular environment [56, 57]. In addition, they can influence myelin maintenance pathways, which rely on lipid synthesis and degradation. Acting as an extracellular sensor, Olr1583 may facilitate metabolic adaptations and reallocation required for myelin recycling and maintenance [58, 59]. Furthermore, Ins2 and Esr1 have been found to have supporting roles in these processes. Upregulation of the Ins2 gene suggests an increased activation of insulin signaling pathways, such as the MAPK/ERK cascade. These pathways are known to promote autophagy and survival under cellular stress. [60]. In an accompanying role, it is well established that increases in estrogen-related pathways play critical roles in cellular homeostasis, particularly supporting autophagy [61, 62]. Taken together, the combined down regulation of Maco1 and Mag, along with upregulation of Olr1583, Ins2, and Esr1 support the consensus that serum-free media in the absence of H2O2 increases transcription of genes associated with a phenotypic and metabolic shift toward autophagy or cellular stress, when compared to serum-containing media.

### 4.9 Chemically Defined Media facilitates pro-regenerative states and cell metabolism in the presence of Deferoxamine Mesylate and 62.5mM H2O2

When both serum-free, CDM, and serum-containing media, D10s, were treated with DFO in the presence of 62.5 mM H2O2, the transcriptional landscape was largely changed (fig 8). The most prominent fold changes were observed in Defb1 (21.09) and Tnfrsf11b (−17.64). Beta defensins, such as Defb1, are known to directly participate in redox-dependent mechanisms. Their reduced form improves anti-microbial function [63]. In SCs, increased ROS levels could induce Defb1 expression to mitigate oxidative stress. The observed upregulation suggests an active oxidative stress management response, which can result from increased ROS production during high metabolic activity or in the presence of H2O2 [64]. Conventionally, an increase oxidative stress would concurrently show a rise in Tnfrsf11b expression. This upregulation can act as a “gauge” of cellular stress, as elevated Tnfrsf11b exacerbates ROS production by mitochondria [65]. If oxidative damage begins to overwhelm these compensatory mechanisms, Tnfrsf11b expression would further increase, ultimately shifting the SC from autophagy toward an apoptotic phenotype [66]. Combined, both the upregulation of Defb1 and downregulation Tnfrsf11b suggest the SCs are actively managing oxidative stress, carefully balancing autophagy and apoptosis. Supporting evidence includes the suppression of Bcl2l11 and Bcl1l12, which is consistent with the assumption that the SCs are indeed successfully regulating oxidative stress [67, 68, 69]. Additionally, changes in Hspbap1 (11.68), Sycp2 (7.45), Ins2 (6.5), Birc3 (−5.42), and Htt (4.64) further indicate a pro-autophagy phenotype, favoring intracellular stability [70, 71, 72, 73]. Notably, Ins2 upregulation suggests an adaptation needed to meet the increased energy demands of these metabolic changes [74], while upregulation of Htt indicates a persisting myelinating phenotype [75]. Overall, the observed transcriptional changes when comparing serum-free vs serum-containing media indicate that the former promotes autophagy and cellular recycling at greater levels, when in the presence of 62.5mM H2O2 and DFO. This adaptation permits the continued repair and maintenance required for SCs to retain their pro-repair and regenerative phenotypes in the more hostile microenvironment of injury.

**Figure 8.**
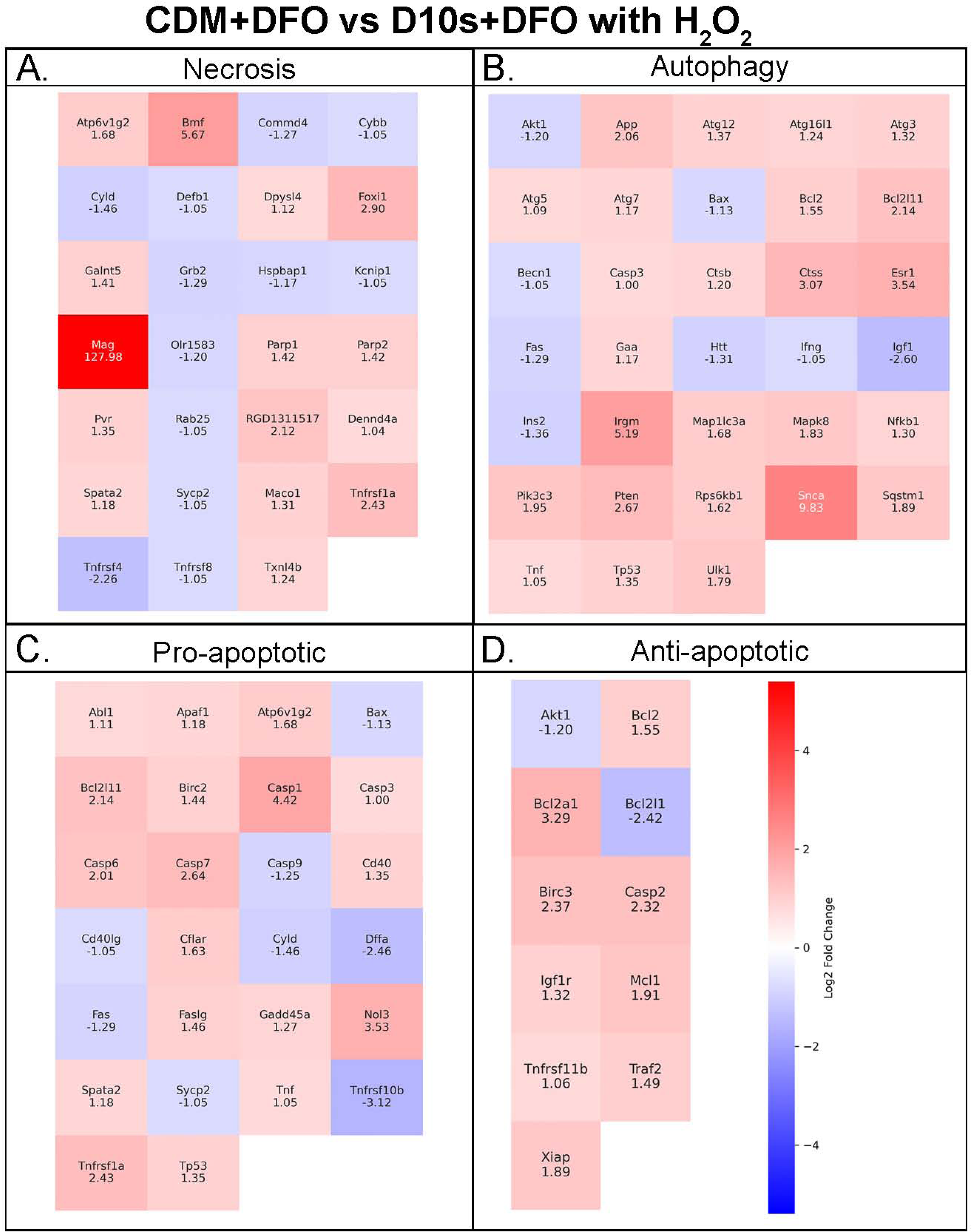
RT-qPCR shows pro-repair and pro-regenerative Schwann Cells in CDM media in the presence of DFO and H2O2. Heatmaps showing Schwann Cell, in serum-free versus serum medium with the presence of DFO and H2O2, transcripts expression associated in **(A)** necrosis, **(B)** autophagy, **(C)** pro-apoptotic, and **(D)** antiapoptotic pathways. Fold-change values shown on heatmap.

## Contributions

**Giles W. Plant, Yee H. E. Ma, Abhinay R. Putta**: Designed the research. **Yee H. E. Ma, Abhinay R. Putta, Cyrus Chan, and Giles W. Plant**: Performed the experiments. **Yee H. E. Ma, Abhinay R. Putta, Cyrus Chan, Stephen S. Vidman:** Imaging **Yee H. E. Ma, Abhinay R. Putta, Cyrus H. H. Chan, Stephen R. Vidman:** Processed raw data using image J. **Cyrus H.H. Chan, Abhinay R. Putta, Stephen R. Vidman and Yee H. E. Ma:** Analyzed the data. **Giles W. Plant, Yee H. E. Ma, Cyrus H. H. Chan, Stephen R. Vidman, Abhinay R. Putta and Paula V. Monje:** Wrote and edited the manuscript.

