## Supplemental Tables for "Efficacy of Deferoxamine Mesylate in Serum and Serum-Free Media: Adult Schwann Cell Survival Following Hydrogen Peroxide Induced Cell Death"

Supplementals

Supplemental Table 1 – Media Components

| Media Abbreviations | Ingredients |
|---|---|
| D10s | 90% DMEM (Dulbecco's Modified Eagle Medium, Gibco), 10% HIFBS (Heat Inactivated Fetal Bovine Serum, Gibco, Aliquoted then heated in 56°C water bath for 30 minutes), Gentamicin (10mg/mL, Gibco), Glutamax (200mM, Gibco) |
| CDM | 100% DMEM/F12 (Dulbecco's Modified Eagle Medium/Nutrient Mixture F-12), Glutamax (200mM, Gibco), Gentamicin (50mg/mL), Bovine Insulin (10µg/mL, Millipore Sigma I6634-100MG), Human Transferrin (10µg/mL, Millipore Sigma T8158-1G), Putrescine dihydrochloride (200µM, Millipore Sigma P5780-5G), Sodium Selenite (30nM, Millipore Sigma S5261-25G) |
| ____ +Mit | Base media plus: Forskolin (2mM, Sigma Aldrich), BPEX (Bovine Pituitary Extract, 20mg/mL) |
| ____ +3F | Base media plus: Forskolin (2mM, Sigma Aldrich), Neuregulin Beta 1 (10ng/mL), BPEX (Bovine Pituitary Extract, 20mg/mL) |
| ____ +DFO | Base media (+/- Mit or +/- 3F) plus: Deferoxamine Mesylate |
| ____ +bFGF | Base media (+/- Mit or +/- 3F) plus: Basic Fibroblast Growth Factor |
| ____ +FGF5 | Base media (+/- Mit or +/- 3F) plus: Fibroblast Growth Factor 5 |

Supplemental Table 2 – Pretreatment Groups for Survival Quantification (Fig. 2, 3, 4)

| Serum Containing (Fig. 3) | Serum-Free (Fig. 4) | Growth Factor & Neurotrophins (Fig. 2) |
|---|---|---|
| CDM -$H_2O_2$ | D10S -$H_2O_2$ | D10S Mit -$H_2O_2$ |
| CDM +$H_2O_2$ | D10S +$H_2O_2$ | D10S Mit +$H_2O_2$ |
| CDM DFO -$H_2O_2$ | D10S DFO -$H_2O_2$ | D10S 3F -$H_2O_2$ |
| CDM DFO +$H_2O_2$ | D10S DFO +$H_2O_2$ | D10s 3F +$H_2O_2$ |
| CDM 3F -$H_2O_2$ | D10S 3F -$H_2O_2$ | D10S Mit bFGF -$H_2O_2$ |
| CDM 3F +$H_2O_2$ | D10S 3F +$H_2O_2$ | D10S Mit bFGF +$H_2O_2$ |
| CDM 3F DFO -$H_2O_2$ | D10S 3F DFO -$H_2O_2$ | D10S Mit FGF5 -$H_2O_2$ |
| CDM 3F DFO +$H_2O_2$ | D10S 3F DFO +$H_2O_2$ | D10S Mit FGF5 +$H_2O_2$ |

Supplemental Table 3 – Pretreatment Groups for RT-qPCR (Fig. 7, 8)

| Serum Containing | Serum-Free |
|---|---|
| CDM -$H_2O_2$ | D10S -$H_2O_2$ |
| CDM DFO +$H_2O_2$ | D10S DFO +$H_2O_2$ |
| CDM 3F +$H_2O_2$ | D10S 3F +$H_2O_2$ |
| CDM 3F DFO +$H_2O_2$ | D10S 3F DFO +$H_2O_2$ |

Supplemental Table 4 – Gene list from RT-qPCR

| | |
|---|---|
| Anti-Apoptotic | Akt1, Bcl2, Bcl2a1 (Bfl-1, A1), Bcl2l1 (Bcl-xl), Bcl2l1 (Bcl-xl), Bcl2l1 (Bcl-xl), Birc3 (c-IAP1), Casp2, Igf1r, Mcl1, Tnfrsf11b (Opg), Traf2, Xiap |
| Pro-Apoptotic | Abl1, Apaf1, Atp6v1g2, Bax, Bcl2l11, Birc2 (c-IAP2), Casp1 (Ice), Casp3, Casp6, Casp7, Casp9, Cd40, Cd40lg, Cflar (Casper), Cyld, Dffa, Fas, Faslg, Gadd45a, Nol3, Spata2, Sycp2, Tnf, Tnfrsf1a (Tnfr1), Tnfrsf10b (QuantiNova Symbol: AABR07018323.1), Tp53 (p53) |
| Necrotic | Atp6v1g2, Bmf, Commd4, Cybb, Cyld, Defb1, Dpysl4, Foxi1, Galnt5, Grb2, Hspbap1, Kcnip1, Mag, Olr1583, Parp1 (Adprt1), Parp2, Pvr, Rab25, RGD1311517, Dennd4a, Spata2, Sycp2, Maco1, Tnfrsf1a (Tnfr1), Tnfrsf4 (Ox40), Tnfrsf8, Txnl4b |
| Autophagy | Akt1, App, Atg12, Atg16l1, Atg3, Atg5, Atg7, Bax, Bcl2, Bcl2l1 (Bcl-xl), Bcl2l1 (Bcl-xl), Bcl2l1 (Bcl-xl), Becn1, Casp3, Ctsb, Ctss, Esr1 (Erα), Fas, Gaa, Htt, Ifng, Igf1, Ins2, Irgm, Map1lc3a, Mapk8 (Jnk1), Nfkb1, Pik3c3 (Vps34), Pten, Rps6kb1, Snca, Sqstm1, TnfTp53 (p53), Ulk1 |

Supplemental Table 5 – P values for cell survival counts in D10S Groups

| | | | |
|---|---|---|---|
| D10S (+) $H_2O_2$ vs D10S (-) $H_2O_2$ | $p$ = 1.157e-13 | D10S 3F DFO (-) $H_2O_2$ vs D10S 3F (-) $H_2O_2$ | $p$ = 1.000e+00 |
| D10S 3F (-) $H_2O_2$ vs D10S (-) $H_2O_2$ | $p$ = 9.999e-01 | D10S 3F DFO (+) $H_2O_2$ vs D10S 3F (-) $H_2O_2$ | $p$ = 4.464e-03 |
| D10S 3F (+) $H_2O_2$ vs D10S (-) $H_2O_2$ | $p$ = 4.063e-14 | D10S DFO (-) $H_2O_2$ vs D10S 3F (-) $H_2O_2$ | $p$ = 5.484e-01 |
| D10S 3F DFO (-) $H_2O_2$ vs D10S (-) $H_2O_2$ | $p$ = 1.000e+00 | D10S DFO (+) $H_2O_2$ vs D10S 3F (-) $H_2O_2$ | $p$ = 5.124e-02 |
| D10S 3F DFO (+) $H_2O_2$ vs D10S (-) $H_2O_2$ | $p$ = 1.747e-02 | D10S 3F DFO (-) $H_2O_2$ vs D10S 3F (+) $H_2O_2$ | $p$ = 3.797e-14 |
| D10S DFO (-) $H_2O_2$ vs D10S (-) $H_2O_2$ | $p$ = 7.974e-01 | D10S 3F DFO (+) $H_2O_2$ vs D10S 3F (+) $H_2O_2$ | $p$ = 1.181e-13 |
| D10S DFO (+) $H_2O_2$ vs D10S (-) $H_2O_2$ | $p$ = 1.455e-01 | D10S DFO (-) $H_2O_2$ vs D10S 3F (+) $H_2O_2$ | $p$ = 8.771e-14 |
| D10S 3F (-) $H_2O_2$ vs D10S (+) $H_2O_2$ | $p$ = 1.039e-13 | D10S DFO (+) $H_2O_2$ vs D10S 3F (+) $H_2O_2$ | $p$ = 9.259e-14 |
| D10S 3F (+) $H_2O_2$ vs D10S (+) $H_2O_2$ | $p$ = 9.664e-02 | D10S 3F DFO (+) $H_2O_2$ vs D10S 3F DFO (-) $H_2O_2$ | $p$ = 7.813e-03 |
| D10S 3F DFO (-) $H_2O_2$ vs D10S (+) $H_2O_2$ | $p$ = 1.090e-13 | D10S DFO (-) $H_2O_2$ vs D10S 3F DFO (-) $H_2O_2$ | $p$ = 6.532e-01 |
| D10S 3F DFO (+) $H_2O_2$ vs D10S (+) $H_2O_2$ | $p$ = 1.577e-08 | D10S DFO (+) $H_2O_2$ vs D10S 3F DFO (-) $H_2O_2$ | $p$ = 7.942e-02 |
| D10S DFO (-) $H_2O_2$ vs D10S (+) $H_2O_2$ | $p$ = 2.240e-11 | D10S DFO (-) $H_2O_2$ vs D10S 3F DFO (+) $H_2O_2$ | $p$ = 5.976e-01 |
| D10S DFO (+) $H_2O_2$ vs D10S (+) $H_2O_2$ | $p$ = 2.397e-10 | D10S DFO (+) $H_2O_2$ vs D10S 3F DFO (+) $H_2O_2$ | $p$ = 9.893e-01 |
| D10S 3F (+) $H_2O_2$ vs D10S 3F (-) $H_2O_2$ | $p$ = 3.730e-14 | D10S DFO (+) $H_2O_2$ vs D10S DFO (-) $H_2O_2$ | $p$ = 9.663e-01 |

Supplemental Table 5 – P values for cell survival counts in CDM Groups

| | | | |
|---|---|---|---|
| CDM (+) $H_2O_2$ vs CDM (-) $H_2O_2$ | $p$ = 1.128e-01 | CDM 3F DFO (-) $H_2O_2$ vs CDM 3F (-) $H_2O_2$ | $p$ = 6.121e-01 |
| CDM 3F (-) $H_2O_2$ vs CDM (-) $H_2O_2$ | $p$ = 9.951e-01 | CDM 3F DFO (+) $H_2O_2$ vs CDM 3F (-) $H_2O_2$ | $p$ = 9.987e-01 |
| CDM 3F (+) $H_2O_2$ vs CDM (-) $H_2O_2$ | $p$ = 4.214e-03 | CDM DFO (-) $H_2O_2$ vs CDM 3F (-) $H_2O_2$ | $p$ = 1.000e+00 |
| CDM 3F DFO (-) $H_2O_2$ vs CDM (-) $H_2O_2$ | $p$ = 1.813e-01 | CDM DFO (+) $H_2O_2$ vs CDM 3F (-) $H_2O_2$ | $p$ = 9.995e-01 |
| CDM 3F DFO (+) $H_2O_2$ vs CDM (-) $H_2O_2$ | $p$ = 8.513e-01 | CDM 3F DFO (-) $H_2O_2$ vs CDM 3F (+) $H_2O_2$ | $p$ = 9.434e-01 |
| CDM DFO (-) $H_2O_2$ vs CDM (-) $H_2O_2$ | $p$ = 9.917e-01 | CDM 3F DFO (+) $H_2O_2$ vs CDM 3F (+) $H_2O_2$ | $p$ = 1.396e-01 |
| CDM DFO (+) $H_2O_2$ vs CDM (-) $H_2O_2$ | $p$ = 1.000e+00 | CDM DFO (-) $H_2O_2$ vs CDM 3F (+) $H_2O_2$ | $p$ = 6.144e-02 |
| CDM 3F (-) $H_2O_2$ vs CDM (+) $H_2O_2$ | $p$ = 5.049e-01 | CDM DFO (+) $H_2O_2$ vs CDM 3F (+) $H_2O_2$ | $p$ = 4.145e-03 |
| CDM 3F (+) $H_2O_2$ vs CDM (+) $H_2O_2$ | $p$ = 9.281e-01 | CDM 3F DFO (+) $H_2O_2$ vs CDM 3F DFO (-) $H_2O_2$ | $p$ = 8.843e-01 |
| CDM 3F DFO (-) $H_2O_2$ vs CDM (+) $H_2O_2$ | $p$ = 1.000e+00 | CDM DFO (-) $H_2O_2$ vs CDM 3F DFO (-) $H_2O_2$ | $p$ = 6.582e-01 |
| CDM 3F DFO (+) $H_2O_2$ vs CDM (+) $H_2O_2$ | $p$ = 8.185e-01 | CDM DFO (+) $H_2O_2$ vs CDM 3F DFO (-) $H_2O_2$ | $p$ = 2.260e-01 |
| CDM DFO (-) $H_2O_2$ vs CDM (+) $H_2O_2$ | $p$ = 5.548e-01 | CDM DFO (-) $H_2O_2$ vs CDM 3F DFO (+) $H_2O_2$ | $p$ = 9.995e-01 |
| CDM DFO (+) $H_2O_2$ vs CDM (+) $H_2O_2$ | $p$ = 1.374e-01 | CDM DFO (+) $H_2O_2$ vs CDM 3F DFO (+) $H_2O_2$ | $p$ = 9.259e-01 |
| CDM 3F (+) $H_2O_2$ vs CDM 3F (-) $H_2O_2$ | $p$ = 5.037e-02 | CDM DFO (+) $H_2O_2$ vs CDM DFO (-) $H_2O_2$ | $p$ = 9.989e-01 |

Supplemental Table 6 – P values for WC-CTCF (Hif1a)

| | | | |
|---|---|---|---|
| CDM (-) $H_2O_2$ vs CDM (+) $H_2O_2$ | *p* = 5.600e-01 | D10S (-) $H_2O_2$ vs D10S (+) $H_2O_2$ | *p* = 1.800e-02 |
| CDM (-) $H_2O_2$ vs CDM 3F (-) $H_2O_2$ | *p* = 1.000e+00 | D10S (-) $H_2O_2$ vs D10S 3F (-) $H_2O_2$ | *p* = 4.000e-02 |
| CDM (-) $H_2O_2$ vs CDM 3F (+) $H_2O_2$ | *p* = 9.890e-01 | D10S (-) $H_2O_2$ vs D10S 3F (+) $H_2O_2$ | *p* = 1.500e-02 |
| CDM (-) $H_2O_2$ vs CDM 3F DFO (-) $H_2O_2$ | *p* = 8.100e-01 | D10S (-) $H_2O_2$ vs D10S 3F DFO (-) $H_2O_2$ | *p* = 1.300e-02 |
| CDM (-) $H_2O_2$ vs CDM 3F DFO (+) $H_2O_2$ | *p* = 9.770e-01 | D10S (-) $H_2O_2$ vs D10S 3F DFO (+) $H_2O_2$ | *p* = 2.900e-02 |
| CDM (-) $H_2O_2$ vs CDM DFO (-) $H_2O_2$ | *p* = 9.390e-01 | D10S (-) $H_2O_2$ vs D10S DFO (-) $H_2O_2$ | *p* = 2.500e-02 |
| CDM (-) $H_2O_2$ vs CDM DFO (+) $H_2O_2$ | *p* = 9.220e-01 | D10S (-) $H_2O_2$ vs D10S DFO (+) $H_2O_2$ | *p* = 2.700e-02 |
| CDM (+) $H_2O_2$ vs CDM 3F (-) $H_2O_2$ | *p* = 1.060e-01 | D10S (+) $H_2O_2$ vs D10S 3F (-) $H_2O_2$ | *p* = 7.000e-05 |
| CDM (+) $H_2O_2$ vs CDM 3F (+) $H_2O_2$ | *p* = 1.280e-01 | D10S (+) $H_2O_2$ vs D10S 3F (+) $H_2O_2$ | *p* = 9.090e-01 |
| CDM (+) $H_2O_2$ vs CDM 3F DFO (-) $H_2O_2$ | *p* = 9.910e-01 | D10S (+) $H_2O_2$ vs D10S 3F DFO (-) $H_2O_2$ | *p* = 8.280e-01 |
| CDM (+) $H_2O_2$ vs CDM 3F DFO (+) $H_2O_2$ | *p* = 7.240e-01 | D10S (+) $H_2O_2$ vs D10S 3F DFO (+) $H_2O_2$ | *p* = 1.000e-03 |
| CDM (+) $H_2O_2$ vs CDM DFO (-) $H_2O_2$ | *p* = 6.370e-01 | D10S (+) $H_2O_2$ vs D10S DFO (-) $H_2O_2$ | *p* = 1.700e-02 |
| CDM (+) $H_2O_2$ vs CDM DFO (+) $H_2O_2$ | *p* = 9.560e-01 | D10S (+) $H_2O_2$ vs D10S DFO (+) $H_2O_2$ | *p* = 3.000e-03 |
| CDM 3F (-) $H_2O_2$ vs CDM 3F (+) $H_2O_2$ | *p* = 9.850e-01 | D10S 3F (-) $H_2O_2$ vs D10S 3F (+) $H_2O_2$ | *p* = 2.300e-04 |
| CDM 3F (-) $H_2O_2$ vs CDM 3F DFO (-) $H_2O_2$ | *p* = 2.630e-01 | D10S 3F (-) $H_2O_2$ vs D10S 3F DFO (-) $H_2O_2$ | *p* = 2.740e-01 |
| CDM 3F (-) $H_2O_2$ vs CDM 3F DFO (+) $H_2O_2$ | *p* = 4.800e-01 | D10S 3F (-) $H_2O_2$ vs D10S 3F DFO (+) $H_2O_2$ | *p* = 1.600e-02 |
| CDM 3F (-) $H_2O_2$ vs CDM DFO (-) $H_2O_2$ | *p* = 1.040e-01 | D10S 3F (-) $H_2O_2$ vs D10S DFO (-) $H_2O_2$ | *p* = 3.000e-03 |
| CDM 3F (-) $H_2O_2$ vs CDM DFO (+) $H_2O_2$ | *p* = 4.440e-01 | D10S 3F (-) $H_2O_2$ vs D10S DFO (+) $H_2O_2$ | *p* = 5.000e-03 |
| CDM 3F (+) $H_2O_2$ vs CDM 3F DFO (-) $H_2O_2$ | *p* = 2.770e-01 | D10S 3F (+) $H_2O_2$ vs D10S 3F DFO (-) $H_2O_2$ | *p* = 5.960e-01 |
| CDM 3F (+) $H_2O_2$ vs CDM 3F DFO (+) $H_2O_2$ | *p* = 5.270e-01 | D10S 3F (+) $H_2O_2$ vs D10S 3F DFO (+) $H_2O_2$ | *p* = 6.000e-03 |
| CDM 3F (+) $H_2O_2$ vs CDM DFO (-) $H_2O_2$ | *p* = 3.840e-01 | D10S 3F (+) $H_2O_2$ vs D10S DFO (-) $H_2O_2$ | *p* = 3.000e-02 |
| CDM 3F (+) $H_2O_2$ vs CDM DFO (+) $H_2O_2$ | *p* = 4.240e-01 | D10S 3F (+) $H_2O_2$ vs D10S DFO (+) $H_2O_2$ | *p* = 1.000e-02 |
| CDM 3F DFO (-) $H_2O_2$ vs CDM 3F DFO (+) $H_2O_2$ | *p* = 9.770e-01 | D10S 3F DFO (-) $H_2O_2$ vs D10S 3F DFO (+) $H_2O_2$ | *p* = 9.180e-01 |
| CDM 3F DFO (-) $H_2O_2$ vs CDM DFO (-) $H_2O_2$ | *p* = 9.590e-01 | D10S 3F DFO (-) $H_2O_2$ vs D10S DFO (-) $H_2O_2$ | *p* = 1.000e+00 |
| CDM 3F DFO (-) $H_2O_2$ vs CDM DFO (+) $H_2O_2$ | *p* = 1.000e+00 | D10S 3F DFO (-) $H_2O_2$ vs D10S DFO (+) $H_2O_2$ | *p* = 9.880e-01 |
| CDM 3F DFO (+) $H_2O_2$ vs CDM DFO (-) $H_2O_2$ | *p* = 1.000e+00 | D10S 3F DFO (+) $H_2O_2$ vs D10S DFO (-) $H_2O_2$ | *p* = 6.800e-02 |
| CDM 3F DFO (+) $H_2O_2$ vs CDM DFO (+) $H_2O_2$ | *p* = 9.990e-01 | D10S 3F DFO (+) $H_2O_2$ vs D10S DFO (+) $H_2O_2$ | *p* = 7.830e-01 |
| CDM DFO (-) $H_2O_2$ vs CDM DFO (+) $H_2O_2$ | *p* = 1.000e+00 | D10S DFO (-) $H_2O_2$ vs D10S DFO (+) $H_2O_2$ | *p* = 5.970e-01 |

Supplemental Table 7 – P values for WC-CTCF (Collagen IV)

| | | | |
|---|---|---|---|
| CDM (-) $H_2O_2$ vs CDM (+) $H_2O_2$ | *p* = 1.000e-03 | D10S (-) $H_2O_2$ vs D10S (+) $H_2O_2$ | *p* = 9.820e-01 |
| CDM (-) $H_2O_2$ vs CDM 3F (-) $H_2O_2$ | *p* = 1.000e+00 | D10S (-) $H_2O_2$ vs D10S 3F (-) $H_2O_2$ | *p* = 2.900e-04 |
| CDM (-) $H_2O_2$ vs CDM 3F (+) $H_2O_2$ | *p* = 9.870e-01 | D10S (-) $H_2O_2$ vs D10S 3F (+) $H_2O_2$ | *p* = 1.000e+00 |
| CDM (-) $H_2O_2$ vs CDM 3F DFO (-) $H_2O_2$ | *p* = 1.810e-01 | D10S (-) $H_2O_2$ vs D10S 3F DFO (-) $H_2O_2$ | *p* = 7.750e-01 |
| CDM (-) $H_2O_2$ vs CDM 3F DFO (+) $H_2O_2$ | *p* = 1.060e-01 | D10S (-) $H_2O_2$ vs D10S 3F DFO (+) $H_2O_2$ | *p* = 9.420e-01 |
| CDM (-) $H_2O_2$ vs CDM DFO (-) $H_2O_2$ | *p* = 2.100e-02 | D10S (-) $H_2O_2$ vs D10S DFO (-) $H_2O_2$ | *p* = 1.000e+00 |
| CDM (-) $H_2O_2$ vs CDM DFO (+) $H_2O_2$ | *p* = 4.100e-02 | D10S (-) $H_2O_2$ vs D10S DFO (+) $H_2O_2$ | *p* = 9.770e-01 |
| CDM (+) $H_2O_2$ vs CDM 3F (-) $H_2O_2$ | *p* = 2.100e-04 | D10S (+) $H_2O_2$ vs D10S 3F (-) $H_2O_2$ | *p* = 1.000e-05 |
| CDM (+) $H_2O_2$ vs CDM 3F (+) $H_2O_2$ | *p* = 1.100e-04 | D10S (+) $H_2O_2$ vs D10S 3F (+) $H_2O_2$ | *p* = 6.150e-01 |
| CDM (+) $H_2O_2$ vs CDM 3F DFO (-) $H_2O_2$ | *p* = 7.900e-02 | D10S (+) $H_2O_2$ vs D10S 3F DFO (-) $H_2O_2$ | *p* = 9.360e-01 |
| CDM (+) $H_2O_2$ vs CDM 3F DFO (+) $H_2O_2$ | *p* = 3.000e-03 | D10S (+) $H_2O_2$ vs D10S 3F DFO (+) $H_2O_2$ | *p* = 1.630e-01 |
| CDM (+) $H_2O_2$ vs CDM DFO (-) $H_2O_2$ | *p* = 1.400e-02 | D10S (+) $H_2O_2$ vs D10S DFO (-) $H_2O_2$ | *p* = 9.270e-01 |
| CDM (+) $H_2O_2$ vs CDM DFO (+) $H_2O_2$ | *p* = 6.730e-01 | D10S (+) $H_2O_2$ vs D10S DFO (+) $H_2O_2$ | *p* = 1.000e+00 |
| CDM 3F (-) $H_2O_2$ vs CDM 3F (+) $H_2O_2$ | *p* = 9.640e-01 | D10S 3F (-) $H_2O_2$ vs D10S 3F (+) $H_2O_2$ | *p* = 1.000e-05 |
| CDM 3F (-) $H_2O_2$ vs CDM 3F DFO (-) $H_2O_2$ | *p* = 1.420e-01 | D10S 3F (-) $H_2O_2$ vs D10S 3F DFO (-) $H_2O_2$ | *p* = 1.000e-05 |
| CDM 3F (-) $H_2O_2$ vs CDM 3F DFO (+) $H_2O_2$ | *p* = 1.600e-02 | D10S 3F (-) $H_2O_2$ vs D10S 3F DFO (+) $H_2O_2$ | *p* = 2.600e-04 |
| CDM 3F (-) $H_2O_2$ vs CDM DFO (-) $H_2O_2$ | *p* = 9.300e-04 | D10S 3F (-) $H_2O_2$ vs D10S DFO (-) $H_2O_2$ | *p* = 0.000e+00 |
| CDM 3F (-) $H_2O_2$ vs CDM DFO (+) $H_2O_2$ | *p* = 5.100e-02 | D10S 3F (-) $H_2O_2$ vs D10S DFO (+) $H_2O_2$ | *p* = 4.400e-04 |
| CDM 3F (+) $H_2O_2$ vs CDM 3F DFO (-) $H_2O_2$ | *p* = 9.500e-02 | D10S 3F (+) $H_2O_2$ vs D10S 3F DFO (-) $H_2O_2$ | *p* = 1.680e-01 |
| CDM 3F (+) $H_2O_2$ vs CDM 3F DFO (+) $H_2O_2$ | *p* = 1.100e-02 | D10S 3F (+) $H_2O_2$ vs D10S 3F DFO (+) $H_2O_2$ | *p* = 8.690e-01 |

| | | | |
|---|---|---|---|
| CDM 3F (+) $H_2O_2$ vs CDM DFO (-) $H_2O_2$ | $p$ = 1.000e-03 | D10S 3F (+) $H_2O_2$ vs D10S DFO (-) $H_2O_2$ | $p$ = 1.000e+00 |
| CDM 3F (+) $H_2O_2$ vs CDM DFO (+) $H_2O_2$ | $p$ = 3.400e-02 | D10S 3F (+) $H_2O_2$ vs D10S DFO (+) $H_2O_2$ | $p$ = 4.110e-01 |
| CDM 3F DFO (-) $H_2O_2$ vs CDM 3F DFO (+) $H_2O_2$ | $p$ = 9.980e-01 | D10S 3F DFO (-) $H_2O_2$ vs D10S 3F DFO (+) $H_2O_2$ | $p$ = 3.000e-02 |
| CDM 3F DFO (-) $H_2O_2$ vs CDM DFO (-) $H_2O_2$ | $p$ = 9.470e-01 | D10S 3F DFO (-) $H_2O_2$ vs D10S DFO (-) $H_2O_2$ | $p$ = 4.330e-01 |
| CDM 3F DFO (-) $H_2O_2$ vs CDM DFO (+) $H_2O_2$ | $p$ = 6.640e-01 | D10S 3F DFO (-) $H_2O_2$ vs D10S DFO (+) $H_2O_2$ | $p$ = 6.650e-01 |
| CDM 3F DFO (+) $H_2O_2$ vs CDM DFO (-) $H_2O_2$ | $p$ = 2.880e-01 | D10S 3F DFO (+) $H_2O_2$ vs D10S DFO (-) $H_2O_2$ | $p$ = 7.930e-01 |
| CDM 3F DFO (+) $H_2O_2$ vs CDM DFO (+) $H_2O_2$ | $p$ = 3.410e-01 | D10S 3F DFO (+) $H_2O_2$ vs D10S DFO (+) $H_2O_2$ | $p$ = 1.000e-02 |
| CDM DFO (-) $H_2O_2$ vs CDM DFO (+) $H_2O_2$ | $p$ = 8.690e-01 | D10S DFO (-) $H_2O_2$ vs D10S DFO (+) $H_2O_2$ | $p$ = 8.700e-01 |
